# Metabolomics and ^13^C Labelled Glucose Tracing to Identify Carbon Incorporation into Aberrant Cell Membrane Glycans in Cancer

**DOI:** 10.1101/2024.04.08.588353

**Authors:** Alfredo Reyes-Oliveras, Abigail E. Ellis, Ryan D. Sheldon, Brian Haab

**Affiliations:** Van Andel Institute, 333 Bostwick Ave NE, Grand Rapids, MI, 49503

**Keywords:** Glycans, Metabolism, LC-MS, Carbohydrate Quantification, Bioenergetics

## Abstract

Cell membrane glycans contribute to immune recognition, signaling, and cellular adhesion and migration, and altered membrane glycosylation is a feature of cancer cells that contributes to cancer progression. The uptake and metabolism of glucose and other nutrients essential for glycan synthesis could underlie altered membrane glycosylation, but the relationship between shifts in nutrient metabolism and the effects on glycans have not been directly examined. To address this possibility, we created a novel method that combines stable isotope tracing with metabolomics to enable direct observations of glucose allocation to nucleotide sugars and cell-membrane glycans. We compared the glucose allocation to membrane glycans of two pancreatic cancer cell lines that are genetically identical but have differing energy requirements. The 8988-S cells had higher glucose allocation to membrane glycans and intracellular pathways relating to glycan synthesis, but the 8988-T cells had higher glucose uptake and commitment of glucose to non-glycosylation pathways. The cells lines differed in requirements of glucose for energy production, resulting in differences in glucose bioavailability for glycan synthesis. The workflow demonstrated here enables studies on the effects of metabolic shifts on the commitment of media nutrients to cell-membrane glycans. The results support a flux-based regulation of glucose commitment glycosylation and a mode of metabolic control of cell functions such signaling, immune recognition, and adhesion and migration.

## Introduction

The glycosylation of membrane glycoconjugates of protein, lipid, or nucleic acids affects tissue organization, cell surface receptor modulation, cellular immune responses, and cellular adhesion and migration. Cancer cell membrane glycans are typically altered when compared with those observed on non-cancer cells,^1^ and altered cell membrane glycosylation can affect many aspects of cellular behavior including altered cellular adhesion and migration with implications for cancer progression.^3,4^ The mechanisms by which membrane glycans become altered in cancer are not well understood. However, cancer cells have been reported to shift their nutrient preferences and utilization, such as by increased uptake and use of glucose for energy and anabolism.^2,3^ Glucose and other nutrients such as fructose, glutamine, acetate, and nucleotides, are required for cell membrane glycan synthesis and therefore alterations in metabolism and energy usage could significantly affect the composition and structure of those glycans.

Previous studies have shown associations between modulations in metabolism and alterations to cell-surface glycans.^4–6^ For example, cancer cells undergoing epithelial-mesenchymal transition (EMT) increased both their glucose metabolism and the glycosylation of a fibronectin variant involved in cell adhesion and migration.^7^ Activated T cells *in vivo* show increased flow of glucose toward anabolic pathways, accompanied by an increase in nucleotide-sugar metabolites that are used for glycosylation.^8^ Related to this, the further differentiation of activated T cells into T helper cells induces a shift of glucose use towards aerobic glycolysis, potentially constraining glucose availability for N-glycosylation.^4^ These observations taken together suggest that metabolic shifts can manifest as altered patterns of cell membrane glycan synthesis. However, a direct mechanistic connection between metabolic shifts and altered cell membrane glycans has not been made. Previous efforts have focused on amounts of glycans and metabolites at specific time points but have not directly examined glucose flux through intracellular metabolism to eventual incorporation into cell-membrane glycans.

The incorporation of the stable isotopes ^13^C or ^15^N into cellular metabolites such as glucose and glutamine enables the tracking by mass-spectrometry metabolomics of the labeled carbon or nitrogen atoms through their incorporation into downstream molecules such as cell membrane glycans.^9,10^ This approach is widely used in studies of cellular metabolic states,^11–13^ and may provide direct measurements of glucose routing to monosaccharide synthesis and cell membrane glycan production. However, the combination of stable isotope labeling and metabolomics has not previously been reported in the study of cell membrane glycans.

Metabolomics methods are powerful tools for identifying the small molecules into which stable isotope labels are incorporated, but they are not readily applied to the large and complex glycans of cell membrane. Here, we report the development of a method that uses metabolomic analysis of ^13^C labelled monosaccharides of cell-membrane glycans to enable direct observations of glucose allocation to monosaccharide production and subsequently to altered membrane glycosylation. We report for the first time that cancer-associated metabolic shifts directly affect glucose allocation to membrane glycan synthesis.

## Methods

### Cell Culture and Subcellular Fractionation

The PaTu8988-S and PaTu8988-T cell lines (referred to as 8988-S and 8988-T) were purchased from ATCC (Manassas, VA); the OVCAR4-EV and OVCAR4-OE cell lines were provided by Dr. Susan Bellis at The University of Alabama at Birmingham; and the 8988-T GFPT-1 knockout was contributed by Dr. Costas Lyssiotis at the University of Michigan. All cell lines were cultured in RPMI-1640 media supplemented with 5% FBS and Penicillin-Streptomycin in a 75 mm dish until they achieved 80% confluency. The cells were detached from the plate surface using a cell scraper and were resuspended in growth media. The resulting cell suspension was centrifuged at 300 × g for 5 minutes, and the pellets were washed twice with 3mL of Cell Wash Solution from the MemPer kit (Thermofisher 89842) followed by centrifugation at 300 × g for 5 minutes. To extract cytosolic proteins, the cells were permeabilized with 0.75 mL of permeabilization buffer from the MemPer kit and incubated with constant mixing at 4°C for 10 minutes. Subsequently, the cells were centrifuged at 16,000 g for 15 minutes, and the cytosolic fraction in the supernatant was collected and transferred to a new tube. To obtain a membrane fraction containing glycoconjugates, the pellet was mixed with 0.5 ml of solubilization buffer. The resulting cytosolic and membrane fractions were either stored on ice for immediate use or aliquoted and stored at −80°C for future use.

### Western Blot

The cytosolic and membrane fractions were subjected to electrophoresis on a Novex™ WedgeWell™ 4 to 20%, Tris-Glycine, 1.0 mm, Mini Protein Gel (Thermofisher, XP04200BOX) for 1.5 hours at 125 V to separate the proteins. The proteins were then transferred onto a PVDF membrane and probed for cytosolic, membrane, and glycan markers. Specifically, GAPDH (ThermoFisher, # MA5-15738) was detected as the cytosolic marker, E-Cadherin (Cell Signal, #3195) as the membrane marker, and ConA (Vector Laboratories, FL-1001-25) as the N-glycan marker. A GFPT-1 (MA5-31739) antibody was used to confirm the absence of GFPT-1 in the 8988-T GFPT-1 knockout cells.

### Glycan Cleavage by Acid Hydrolysis

To release monosaccharides from the isolated glycans, the samples underwent digestion with either trifluoroacetic acid (TFA) or acetic acid for acid hydrolysis. TFA was used to cleave any glycosidic bond between neutral monosaccharides in the glycans, while acetic acid was used to cleave only the bonds with N-acetyl neuraminic acid. Briefly, 50 µg of protein from the samples were exposed to either 2M TFA or acetic acid for three hours at 100°C or two hours at 80°C, respectively. The resulting samples were then dried using rotatory evaporation and derivatized with PMP.

### Derivatization of Monosaccharides with 1-Phenyl-3-Methyl-5-Pyrazolone (PMP)

To enable the separation of the carbohydrate isomers, PMP was used to derivatize the cleaved glycans and standard solutions of monosaccharides as controls. The mixed standard solutions containing 0.5 mg/mL of glucose, galactose, mannose, N-acetyl-glucosamine, N-acetyl-galactosamine, glucosamine, galactosamine, or L-fucose were prepared in sterile water. 50 µL of a standard solution or a sample-derived monosaccharide solution were mixed with a PMP solution to a final concentration of 0.1 M PMP in 15% ammonia and 50% methanol. The mixtures were incubated at 70 °C for 1 hour, dried by vacuum centrifugation, reconstituted in 500 µL of water, and washed three times with chloroform. The mixtures were transferred to a new tube containing either underivatized standard of N-acetyl neuraminic acid or samples treated with acetic acid to obtain N-acetyl neuraminic acid from glycans.

### 13C Carbon Tracing

The cell lines were cultured in glucose-free RPMI-1640 media supplemented with 5% dialyzed FBS and P/S and 10 mM of ^13^C-U-Glucose (CLM-1396, Cambridge Isotope Laboratories). The cells were collected at 8, 12, 24, 48, and 72 hours, to examine the contribution of glucose towards glycosylation. The collected samples were subjected to membrane isolation, acid hydrolysis, and PMP derivatization to quantify the amount of ^13^C labeled monosaccharides in the glycans. This method allowed for the measurement of glycosylation changes over time, and the data obtained helped to elucidate the dynamics of glycosylation in response to changes in glucose metabolism.

### Liquid Chromatography-Mass Spectrometry

PMP-derivatized standards and samples were analyzed with a Vanquish liquid chromatography system coupled to an Orbitrap ID-X (Thermo Fisher Scientific) using an H-ESI (heated electrospray ionization) source in positive mode. 2 μL of each standard and sample was injected and run through a 10-minute reversed-phase chromatography CORTECS T3 column (1.6 μm, 2.1mm × 150mm, 186008500, Waters, Eschborn, Germany) combined with a VanGuard pre-column (1.6 μm, 2.1 mm × 5 mm, 186008508, Waters). Buffer A consisted of 100% LC/MS grade water (W6-4, Fisher), 0.01% ammonium hydroxide (A470, Fisher), 5mM ammonium acetate (73594, Sigma), and buffer B consisted of 99% LC/MS grade acetonitrile (A955, Fisher). Column temperature was kept at 30 °C, flow rate was held at 0.3 mL/min, and the chromatography gradient was as follows: 0-1 min from 0% B to 20% B, 1-4 min from 20% B to 30% B, 4-7.5 min from 30% B to 45% B, 7.5-8.5 min from 45% B to 100% B, and 8.5-10 min held at 100% B. A 5 minute wash gradient was run between every injection to flush the column and to re-equilibrate solvent conditions as follows: 0-2 min held at 100% B, 2-3 min from 100% B to 0% B, and 3-5 min held at 0% B. Mass spectrometer parameters were: source voltage 3500V, sheath gas 70, aux gas 25, sweep gas 1, ion transfer tube temperature 300°C, and vaporizer temperature 250°C. Full scan data were collected using the orbitrap with a scan range of 105-1200 m/z at a resolution of 120,000. Fragmentation was induced in the orbitrap with assisted higher-energy collisional dissociation (HCD) collision energies at 20, 40, 60, 80, and 100%. The collision-induced dissociation (CID) energy was fixed at 35%, and resolution was set at 30,000. Data were analyzed using Compound Discoverer (version 3.3), FreeStyle (version 1.8), and TraceFinder (version 5.1) from Thermo Fisher. To correct for natural isotope abundances in labeled samples, a set of unlabeled samples was processed using ^12^C-glucose in place of ^13^C-labeled glucose. FluxFix Isotopologue Analysis Tool^14^ (version 0.1) was used to calculate mass isotopologue distributions.

### Glycan Profiling by Flow Cytometry

Glycan-binding proteins, consisting of lectins (Vector Laboratories), antibodies, and the SiaFind engineered protein (Lectenz), were prepared in BD Pharmingen Stain Buffer. A total of 100,000 cells were incubated with 2 µg/mL of FITC-conjugated glycan-binding proteins or FITC-conjugated streptavidin for 1 hour at 4°C. Subsequently, samples were run in BD Accuri C6 at a medium flow rate (35 µL/min), and 5000 events were collected in the singlets gate. The data analysis was carried out in FCS Express and are presented as the geometric mean and standard error/deviation.

### Bioenergetics Analyses

The Seahorse Analysis was conducted using the MitoStress test according to the manufacturer’s instructions. For the determination of nutrient dependence on glucose, L-glutamine, or fatty acids, acute injections of 2 µM UK5099 (mitochondrial pyruvate carrier inhibitor), 3 µM BPTES (glutaminase inhibitor), or 4 µM Etomoxir (carnitine-palmitoyl transferase inhibitor) were administered after six basal measurements. A fourth injection of 30 µM monensin was introduced^15,16^ to assess ATP production from glycolysis and oxidative phosphorylation. A DMSO injection served as the control.

### Statistical Analysis

All data are presented as mean ± SD, and all analyses were performed in GraphPad Prism 10. We used two-tailed Student t-tests to assess differences between two groups. In instances involving multiple groups, we used a two-way ANOVA with Fisher’s LSD test or Bonferroni post-hoc tests.

## Results

### Derivatization and Identification of Monosaccharides by LC-MS

We developed a strategy to quantify the incorporation of glucose-derived ^13^C carbons into the monosaccharides in cell-membrane glycans (Figure 1A). Several of the monosaccharides are structurally similar isomers (Figure 1B) and cannot be distinguished by retention time or mass-to-charge ratio in LC-MS. We therefore derivatized the neutral monosaccharides with 1-phenyl-3-methyl-5-pyrazolone (PMP) (Figure 1C)^17–19^ to enable their resolution in liquid chromatography (Supplementary Figure 1). We used three unique mixtures of monosaccharides containing one monosaccharide from each isomeric group as standards (Table 1) and derivatized each mixture with PMP. N-acetyl neuraminic acid (Neu5Ac) could not be derivatized by PMP due to its negative charge^19,20^ and was added to a mixture after PMP derivatization.

**Figure 1.**
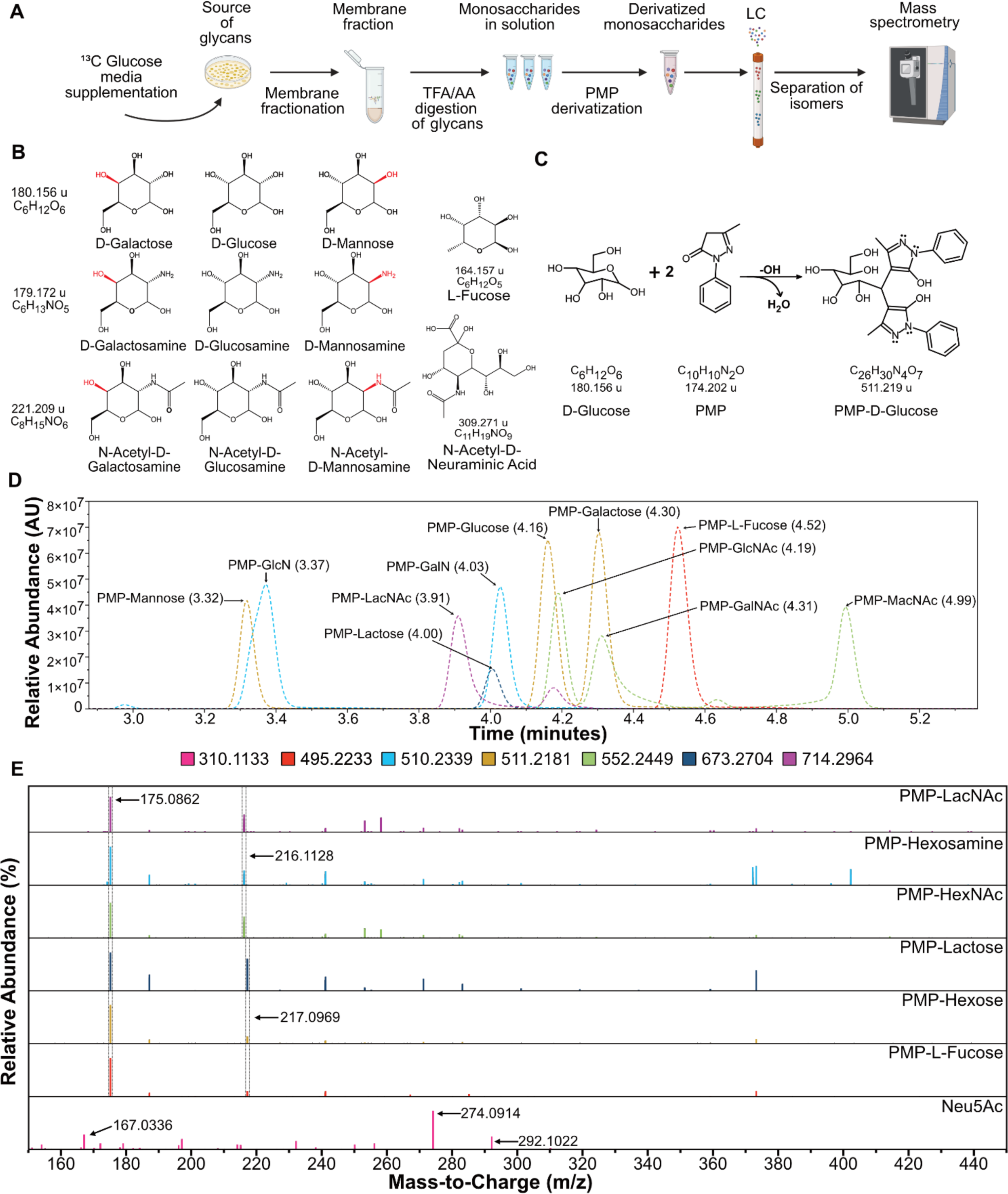
Specific detection of monosaccharide isomers. **A.** Overall strategy for the tracking of glucose from the media to the glycans in cellular membranes. **B.** Structures of monosaccharides found in glycans in humans showing isomer groups, hexoses, hexosamines, N-acetyl-hexosamines, and L-fucose and N-acetyl-neuraminic acid. **C.** The chemical reaction of neutral carbohydrates with PMP to form derivatized PMP-carbohydrates. **D.** Chromatogram showing retention peaks of standard solutions of monosaccharides derivatized with PMP. **E.** MS^2^ spectra of all detected carbohydrates.

**Table 1.** Mono- and di-saccharides used as standards. N-acetyl-glucosamine (GlcNAc), N-acetyl-galactosamine (GalNAc), N-acetyl-mannosamine (ManNAc), glucosamine (GlcN), galactosamine (GalN),), N-acetyl-neuraminic acid (Neu5Ac),), N-acetyl-lactosamine (LacNAc)

| Name | Chemical Formula | PMP Derivatized Formula | Retention Time | Parent Ion [M+H] | Ion Fragments [M+H] |
| --- | --- | --- | --- | --- | --- |
| **GlcNAc** | C<sub>8</sub>H<sub>15</sub>NO<sub>6</sub> | C<sub>28</sub>H<sub>33</sub>N<sub>5</sub>O<sub>7</sub> | 4.19 | 552.2453 | 175.0862, 216.1128 |
| **GalNAc** | C<sub>8</sub>H<sub>15</sub>NO<sub>6</sub> | C<sub>28</sub>H<sub>33</sub>N<sub>5</sub>O<sub>7</sub> | 4.31 | 552.2453 | 175.0862, 216.1128 |
| **MacNAc** | C<sub>8</sub>H<sub>15</sub>NO<sub>6</sub> | C<sub>28</sub>H<sub>33</sub>N<sub>5</sub>O<sub>7</sub> | 4.99 | 552.2453 | 175.0862, 216.1128 |
| **Mannose** | C<sub>6</sub>H<sub>12</sub>O<sub>6</sub> | C<sub>26</sub>H<sub>30</sub>N<sub>4</sub>O<sub>7</sub> | 3.32 | 511.2187 | 175.0862, 217.0969 |
| **Glucose** | C<sub>6</sub>H<sub>12</sub>O<sub>6</sub> | C<sub>26</sub>H<sub>30</sub>N<sub>4</sub>O<sub>7</sub> | 4.16 | 511.2187 | 175.0862, 217.0969 |
| **Galactose** | C<sub>6</sub>H<sub>12</sub>O<sub>6</sub> | C<sub>26</sub>H<sub>30</sub>N<sub>4</sub>O<sub>7</sub> | 4.3 | 511.2187 | 175.0862, 217.0969 |
| **GlcN** | C<sub>6</sub>H<sub>13</sub>NO<sub>5</sub> | C<sub>26</sub>H<sub>31</sub>N<sub>5</sub>O<sub>6</sub> | 3.37 | 510.2347 | 175.0862, 216.1128 |
| **GalN** | C<sub>6</sub>H<sub>13</sub>NO<sub>5</sub> | C<sub>26</sub>H<sub>31</sub>N<sub>5</sub>O<sub>6</sub> | 4.03 | 510.2347 | 175.0862, 216.1128 |
| **L-Fucose** | C<sub>26</sub>H<sub>30</sub>N<sub>4</sub>O<sub>7</sub> | C<sub>26</sub>H<sub>30</sub>N<sub>4</sub>O<sub>6</sub> | 4.52 | 495.2238 | 175.0862, 217.0969 |
| **Neu5Ac** | C<sub>11</sub>H<sub>19</sub>NO<sub>9</sub> | - | 1.07 | 310.1133 | 292.1022, 274.0914, 167.0336 |
| **Lactose** | C<sub>12</sub>H<sub>22</sub>O<sub>11</sub> | C<sub>32</sub>H<sub>40</sub>O<sub>12</sub>N<sub>4</sub> | 4.00 | 673.2715 | 175.0862, 217.0969 |
| **LacNAc** | C<sub>14</sub>H<sub>25</sub>NO<sub>11</sub> | C<sub>34</sub>H<sub>43</sub>N<sub>5</sub>O<sub>12</sub> | 3.91/4.18 | 717.2981 | 175.0862, 216.1128 |

This approach enabled rapid (10-minute) LC-MS separations of all PMP-derivatized and underivatized carbohydrates (Figure 1D). The mass spectra of the parent ions (Supplementary Figure 1) and MS^2^ fragmentation spectra (Figure 1E), in conjunction with unique retention times (Figure 1D), allowed the identification of each carbohydrate in each isomeric grouping. The peak at 175.0862 m/z is the common PMP fragment from all derivatized PMP-sugars. For all amine-containing carbohydrates (e.g., hexosamines and N-acetyl-hexosamines), the second most abundant fragment ion was the peak at 216.1128 m/z, and for the other carbohydrates except for Neu5Ac, the peak at 217.0969 m/z. These peaks arise from the cleavage of C2-C3 and C1-PMP bonds (Supplementary Figure 1). For Neu5Ac, we detected peaks at 292.1022, 274.0914, and 167.0336 m/z, coming from the loss of one or two molecules of H_2_O and cross-ring cleavage, respectively (Supplementary Figure 1C). These masses were further confirmed in complete, high-resolution MS^2^ spectra for each PMP-carbohydrate using collision-induced dissociation (CID) fragmentation (Supplementary Figure 2),^19^ demonstrating unambiguous identification and chromatographic resolution of each monosaccharide.

### Detection of ^13^C Glucose-Labeled Monosaccharides in Membrane-Bound Glycans

After the separation of membrane fractions from cultured cells by subcellular fractionation, the membrane fractions showed high amounts of N-glycans (detected by concanavalin A) and low amounts of the cytosolic marker GAPDH, in contrast to the cytosolic fractions (Figure 2A). We hydrolyzed the membrane fraction with trifluoroacetic acid (TFA) or acetic acid to cleave the glycosidic bonds between the glycan monosaccharides, after which we derivatized the purified monosaccharides with PMP. LC-MS analysis revealed seven monosaccharides: PMP-glucose, PMP-galactose, PMP-mannose, PMP-L-fucose, PMP-glucosamine, PMP-galactosamine, and Neu5Ac (Figure 2B). Mammalian glycans do not contain hexosamines,^21^ but because reaction with TFA removes the acetyl group in N-acetyl hexosamines leading to the formation of hexosamines,^22,23^ we interpreted the presence of hexosamines as a surrogate for N-acetyl-hexosamines. Thus, these observations confirmed the unique identification of the primary monosaccharide components of membrane-bound glycans.

**Figure 2.**
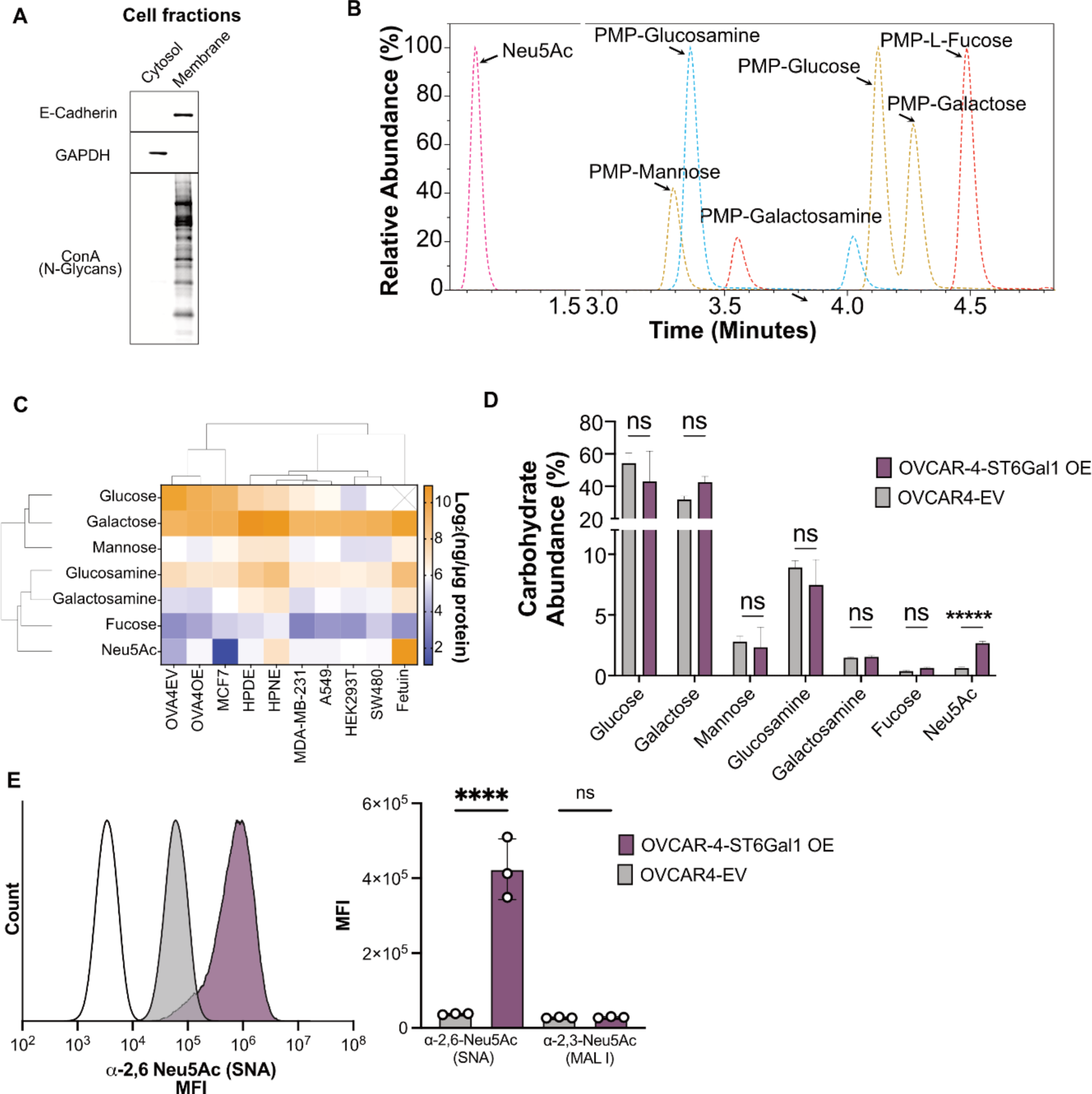
Detection and quantification of cell membrane monosaccharides. **A.** Western Blot showing isolation of cells in cytosolic fraction or membrane fraction. E-Cadherin (membrane marker), GAPDH (cytosolic marker), and Con-A for N-Glycan marker. **B.** Liquid chromatography showing PMP-derivatized carbohydrates isolated from glycans after acid hydrolysis. **C**. Monosaccharide distribution across cell lines and glycoprotein fetuin. **D**. Monosaccharide abundance of OVCAR4-EV and ST6Gal1-OE. **E**. Validation of ST6Gal1 overexpression by flow cytometry using lectins SNA and MAL I. MFI, mean fluorescence intensity. Asterisks with * indicating p < 0.05, ** for p < 0.01, *** for p < 0.001, and **** for p < 0.0001.

We asked whether the distributions of monosaccharides in the cell lines and the purified glycoprotein fetuin corresponded to predicted abundances based on previous studies.^22,23^ We developed external standard curves of the derivatized monosaccharides to enable comparisons of the absolute levels of each monosaccharide (Supplementary Figure 3). The standard curves showed high precision and linearity in the measurements (R^2^ > 0.996) and similar detection limits across the standards, confirming applicability to each of the monosaccharides measured in this study. Comparisons of the absolute levels of monosaccharides (Figure 2C) showed that galactose was the predominant monosaccharide in all samples, which corresponds to the presence of galactose in most types of glycans, including N-glycans, O-glycans, and glycosphingolipids (GSLs), as well as less common proteoglycans like alpha-distro-glycans, collagen-Hyl-Gal, and proteoglycan-O-Xyl.^24^ Fucose and Neu5Ac were the least abundant, consistent with their positions as capping features of glycans.

We tested the accuracy of the method using the OVCAR4 ovarian cancer cell line with overexpression (OE) or empty vector (EV) of the ST6ALl1 glycosyltransferase, which transfers Neu5Ac from CMP-Neu5Ac to galactose in an α-2,6-glycosidic bond. The OE cell line showed a ∼4-fold higher Neu5Ac content than the EV version with no differences in the other monosaccharides (Figure 2D). We confirmed the differences by flow cytometry with two lectins, SNA or MAL I, that respectively bind to α-2,6-Neu5Ac and α-2,3-Neu5Ac. The SNA lectin, but not MAL I, showed significantly increased mean fluorescence intensity (MFI) in the OE cell line. These results confirm unbiased and accurate comparisons of the monosaccharide compositions of cell-membrane glycans.

### Detection of ^13^C Incorporation into Membrane-Bound Glycans

We then tested the ability to detect stable isotopes that were incorporated into the monosaccharides of cell-membrane glycans. We cultured the 8988-T and 8988-S cell lines in media containing ^13^C-glucose (all six carbons with the stable isotope) over three days. Imported glucose undergoes phosphorylation to enter glycolysis and is then processed to form the nucleotide sugars that serve as monosaccharide donors in glycan synthesis (Figure 3A). The phosphorylated glucose can be directed towards various metabolic pathways (Supplementary Figure 4), including the pentose phosphate pathway (PPP) for nucleotide synthesis, the Leloir pathway for UDP-hexose synthesis, or isomerization to fructose-6P. Fructose-6P can undergo further oxidation through glycolysis, pass to the hexosamine biosynthesis pathway (HBP) for nucleotide amino-sugar synthesis, or pass to the fructose-mannose pathway for GDP-hexose synthesis (Figure 3A).

**Figure 3.**
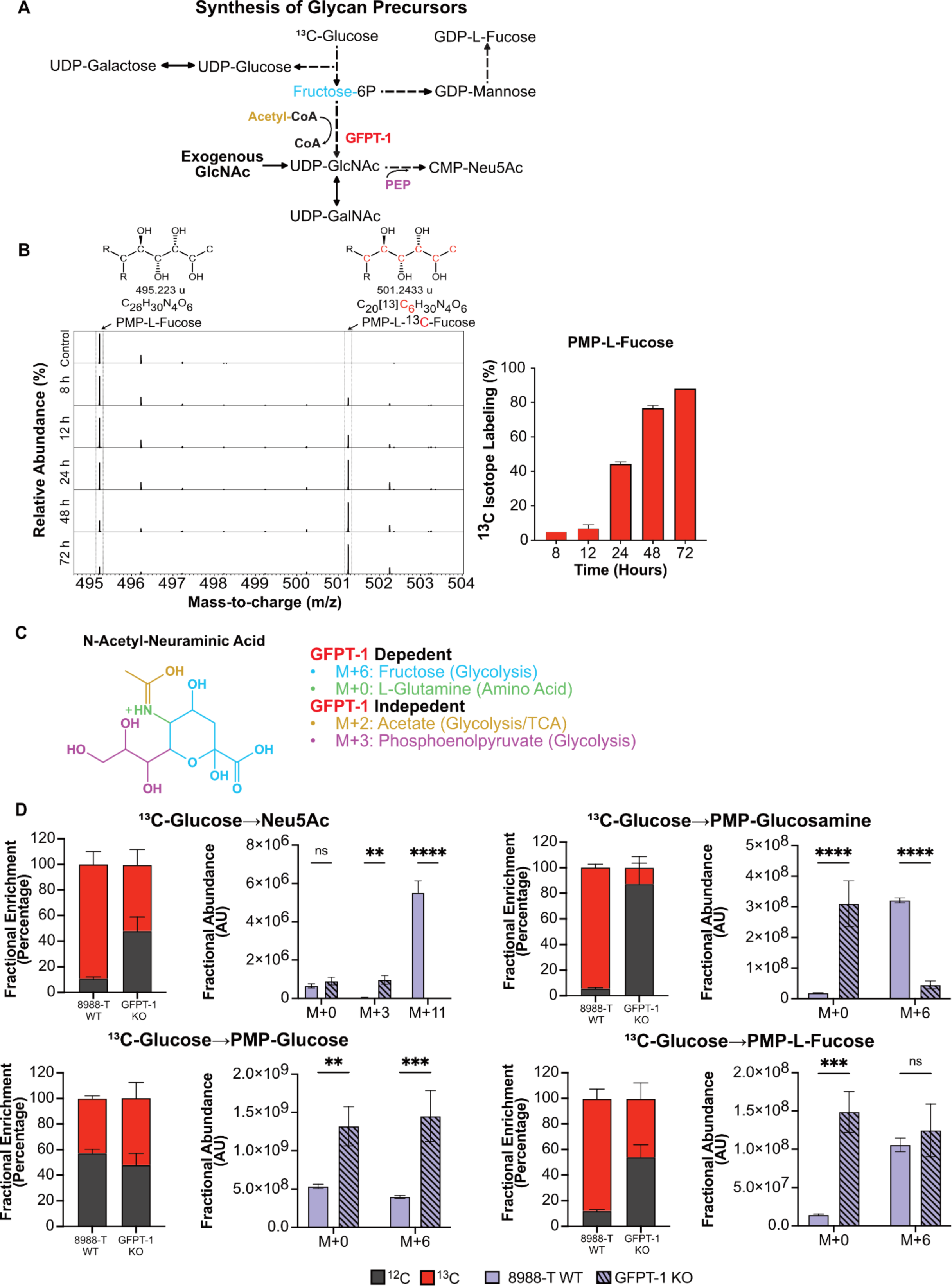
Detection of ^13^C isotope incorporation from ^13^C-glucose to cell-membrane glycans. **A.** Metabolic pathways involved in nucleotide sugar synthesis derived from glycolysis. GFPT-1 in red represents the knockout of the rate-limiting enzyme of the HBP. Phosphoenolpyruvate (PEP). **B.** Mass spectra of PMP-L-Fucose showing ^13^C isotope enrichment after 72 h of cells growing in ^13^C-U-Glucose. R represents the PMP molecule. **C.** Molecular representation of Neu5Ac, providing a visual representation of the metabolites essential for its synthesis and the pathways from which they are derived. **D.** Fractional enrichment and mass isotopologue distribution (MID) showing ^13^C incorporation in derivatized monosaccharides after knockout of GFPT-1. Asterisks with * indicating p < 0.05, ** for p < 0.01, *** for p < 0.001, and **** for p < 0.0001.

We isolated the membrane-bound glycans over several time points and analyzed the isotope abundances in the monosaccharides. The mass shift expected for galactose, mannose, and fucose produced from the incorporation of ^13^C carbons from isotope-labeled glucose is M+6. We observed a consistent increase over 3 days in the M+6 fucose peak at 501.243 m/z, the peak corresponding to PMP-^13^C_6-_fucose, confirming a steady incorporation of ^13^C carbons into L-fucose cleaved from membrane-bound glycoconjugates (Figure 3B).

We then sought to confirm that the ^13^C-labeled glucose used for the synthesis of membrane glycans was processed through the nucleotide sugar synthesis pathways. We compared the monosaccharides obtained from membrane-bound glycoconjugates in the wildtype (WT) 8988-T cell line and in a version with knockout (KO) of the GFPT-1 gene (Supplementary Figure 5).^31^ GFPT-1 catalyzes the rate-limiting step of the hexosamine biosynthesis pathway (HBP) that leads to UDP-GlcNAc synthesis (Figure 3A),^32,33^ the core monosaccharide for N-linked glycosylation. UDP-GlcNAc is also used in the synthesis of UDP-GalNAc and CMP-Neu5Ac, with the latter used in the production of sialic acids. We supplemented the media of the KO cells with GlcNAc due to the growth-inhibitory consequences of GFPT-1 loss.^31^ The expected mass shift of Neu5Ac is M+11, based on M+6 from N-acetyl-mannosamine (an isomer from GlcNAc), M+3 from phosphoenol pyruvate (PEP) arising from glucose metabolism through glycolysis, and M+2 from acetate from glycolysis (Figure 3C). We observed mass shifts in the OK cells consistent with a loss of glucose routing through the HBP (Figure 3D). M+6 ^13^C labeling of PMP-glucosamine and M+11 labeling of N-acetyl-neuraminic acid were significantly reduced in the KO cells relative to the WT cell. M+3 labeling of N-acetyl-neuraminic acid was higher in the GFPT-1 KO cells, consistent with the use of 3 carbons of glucose in the incorporation of PEP into N-acetyl-mannosamine derived from GlcNAc supplementation (Figure 3A and 3C).

Interestingly, we observed an increase in total PMP-L-fucose and PMP-glucose and a decrease in total Neu5Ac in the GFPT-1 knockout cell line (Figure 3D). A decrease in total Neu5Ac was not expected because of the supplementation with GlcNAc to supply the production of UDP-GlcNAc and CMP-Neu5Ac. The change in the abundance of monosaccharides was also observed in PMP-Mannose and PMP-Galactose and was validated by flow cytometry using lectins that bind to epitopes with these monosaccharides (Supplementary Figure 5). These changes suggest that upon loss of GFPT-1 the cells potentially shift glucose toward the Leloir pathway, which produces UDP-glucose and UDP-galactose, and the fructose-mannose pathway, which produces GDP-mannose and GDP-L-fucose (Figure 3A). These findings need further exploration to confirm changes in the intracellular pool of nucleotide sugars. In sum, these results confirm that we are able to detect isotope labels that arise from supplemented ^13^C-glucose and that are processed through glycolysis. Furthermore, we can detect changes in glucose routing from glycolysis to monosaccharide metabolism and glycan synthesis.

### Differential Commitment of Glucose to Membrane Glycans

The ability to track the incorporation of ^13^C carbons from supplemented ^13^C-glucose to membrane glycans allowed us to test whether the commitment of glucose toward monosaccharide metabolism and membrane glycosylation is affected by cancer-associated shifts in glucose uptake or metabolism. We compared glucose routing to membrane glycans between two pancreatic cancer cells lines that are genetically identical, 8988-S and 8988-T, but that have differences in metabolism and glycosylation. The 8988-S cells have a lipogenic phenotype with epithelial-like features, while 8988-T cells have a glycolytic phenotype with mesenchymal features.^34–36^

We monitored ^13^C isotope incorporation from media ^13^C-glucose into the membrane-bound glycoconjugates of the cell lines over 3 days. The 8988-S cells showed higher fractional abundance and total abundance of ^13^C carbon in all monosaccharides except Neu5Ac (Figure 4A). The depletion in ^12^C also was higher in 8988-S, suggesting faster turnover of the membrane glycans. This result supports a higher glucose allocation towards glycan synthesis in the 8988-S cells.

**Figure 4.**
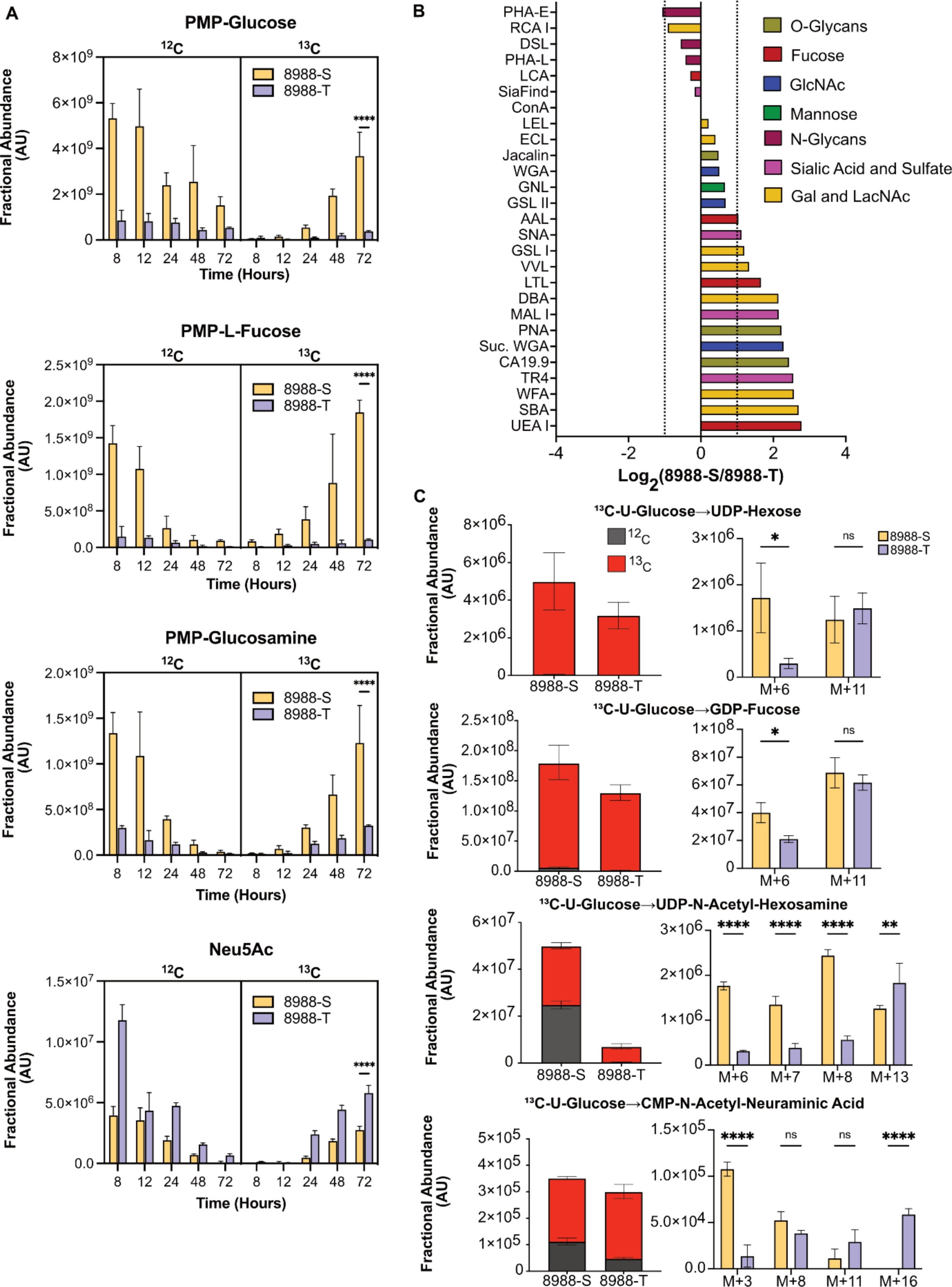
Differential commitment of glucose to membrane glycans. **A.** Fractional abundance of ^13^C and ^12^C across derivatized monosaccharides after labeling with ^13^C glucose for 72 hours. **B**. Fold change analysis of glycosylation between 8988-S and 8988-T. **C.** Fractional abundance of ^13^C and ^12^C and MID of nucleotide sugars. Asterisks with * indicating p < 0.05, ** for p < 0.01, *** for p < 0.001, and **** for p < 0.0001.

We asked whether the greater abundance of monosaccharides in the 8988-S cells was also reflected in the intact cell-surface glycans. We used flow cytometry of 27 lectins and glycan-binding antibodies to determine the relative quantifications of multiple glycan motifs on the cell surfaces (Figure 4B). The 8988-S cells showed higher levels than the 8988-T cells in the targets of most lectins, with 2-4-fold higher levels of fucosylation (AAL, LTL, UEA I) and extensions containing HexNAc and galactose (SBA, WFA, DBA, VVL, GSL I). The greater cell-surface abundances of these glycans in the 8988-S cells are consistent with the increased intracellular glucose commitment to monosaccharide production.

We further investigated the difference between the cell lines in glucose commitment to glycan production by monitoring the ^13^C-glucose labeling in the intracellular pathways related to nucleotide-sugar synthesis (Figure 4C). The total amounts and the ^13^C-labeled amounts of UDP-hexose (representing UDP-glucose and UDP-galactose) and GDP-fucose were substantially higher in the 8988-S cells, mirroring the higher amounts observed in the membrane glycans. The other monosaccharides stemming from the HBP, UDP-GlcNAc, and UDP-GalNAc (Figure 4C) (measured jointly by UDP-HexNAc), also were more abundant in the 8988-S cells, with ∼7-fold higher total amounts and ∼3 fold more ^13^C labeling. The total and ^13^C-labeled CMP-Neu5Ac was similar between the 8988-S cells and the 8988-T cells, whereas cell-membrane Neu5Ac was higher in the 8988-T cells. The difference in cell-membrane levels between 8988-T and 8988-S potentially stem from differential sialyltransferase activity. Nevertheless, this result confirms that 8988-T cells maintain relatively higher Neu5Ac production than the other monosaccharides. In sum, the intracellular levels of the nucleotide sugar precursors generally match the membrane-bound levels of the corresponding monosaccharides. The results also support the conclusion that the 8988-S cells have higher glucose allocation towards monosaccharide metabolism related to nucleotide sugar synthesis and glycan synthesis.

To determine whether the difference in ^13^C incorporation and abundance was from the carbohydrate moiety of the nucleotide sugars or the ribose nucleotide portion, we analyzed the mass isotopologue distribution (MID) of the nucleotide sugars (Figure 4C). The ribose arises from the pentose phosphate pathway (PPP), an offshoot of the second step of glycolysis. The 8988-S cells showed a higher fractional enrichment of the carbohydrate moiety arising from the incorporation of ^13^C carbons coming from the Leloir pathway for UDP-hexose (M+6) and fructose-mannose metabolism for GDP-fucose (M+6). UDP-HexNAc showed higher fractional enrichment in M+6 arising from 6 carbons of glucose through the first step of HBP, M+7 arising from 6 carbons of glucose through the first step of HBP and one carbon allocated in the ring of the uridine and M+8 arising from the HexNAc completely labeled. CMP-Neu5Ac showed enrichment in M+3 arising from PEP synthesis through glycolysis (Figure 3A). The 8988-T cells, on the other hand, showed higher enrichment of isotopes from ribose arising from the PPP combined with the monosaccharide moiety arising from nucleotide sugar synthesis. The enriched isotopologues were M+11 (^13^C_5_-ribose-containing UDP/GDP combining with ^13^C_6_-hexose) in UDP-hexose and GDP-fucose, M+13 (^13^C_5_-ribose-containing UDP combining with ^13^C_8_-N-HexNAc) in UDP-HexNAc, and M+16 (^13^C_5_-ribose-containing CMP combining with ^13^C_11_-Neu5Ac). These results indicates that, although the amount of glucose committed to monosaccharide production through nucleotide sugar synthesis was lower in the 8988-T cells, the glucose flux to the pentose phosphate pathway that produces the ribose for nucleotide synthesis was higher.

### High Energy Demand Limits Glucose Bioavailability for Glycan Synthesis

We asked whether the higher glucose flow through glycolysis and lower glucose flow to glycan production in the 8988-T cells corresponded with higher glucose usage in other non-glycosylation pathways besides the PPP. The 8988-T cells have ∼2-fold higher glucose uptake with similar lactate secretion (Figure 5A), consistent with higher overall glucose usage. In addition, based on Seahorse analysis after acute treatments of the cells with mitochondrial transport inhibitors of pyruvate (UK5099), L-glutamine (BPTES), and fatty acids (etomoxir) (Figure 5B), the 8988-T cells depended primarily on glucose for energy production by oxidative phosphorylation (OXPHOS) using oxygen consumption rate (OCR) as a surrogate, while the 8988-S cells depended primarily on fatty acids (Figure 5C). The cell lines had similar apportionments of ATP production between glycolysis and OXPHOS (Figure 5D), as determined by transformations of the ECAR and OCR measurements (Supplementary Figure 6) to glycolytic and oxidative ATP production rates (J_ATP_).^16^ However, the overall production of ATP was approximately 3 times higher in 8988-T cells than in 8988-S cells (Figure 5D). This higher ATP production by both glycolysis and OXPHOS and the reliance on glucose for mitochondrial ATP production in 8988-T cells is consistent with its 3.2-fold higher proliferation rate (Supplementary Figure 6).

**Figure 5.**
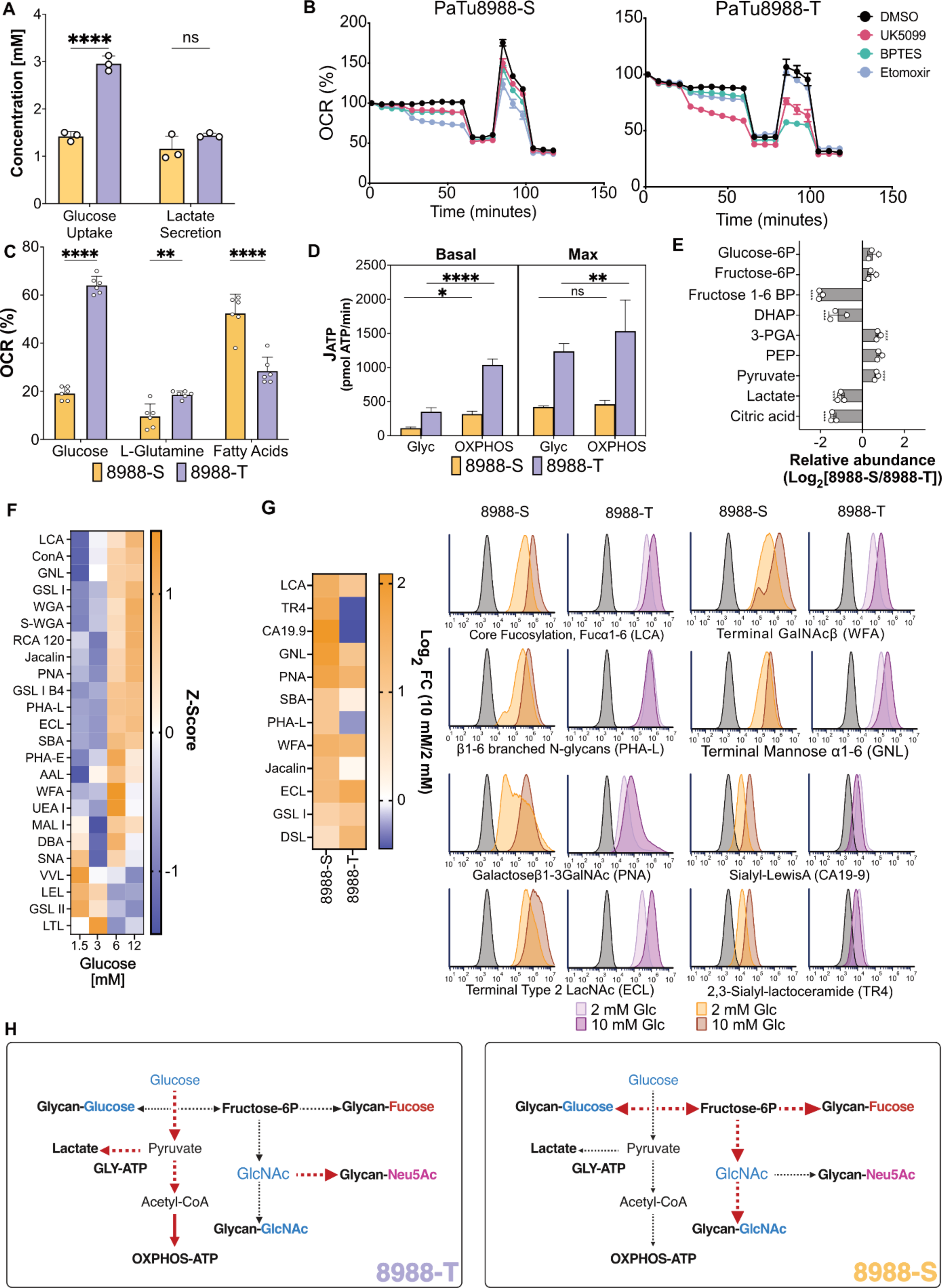
Differential glucose availability constrains glucose commitment to glycosylation. **A.** Measurement of glucose uptake and lactate secretion in 8988-S and 8988-T. **B.** Nutrient dependency measured by Seahorse analysis using inhibitors that target the transporters UK5099 (MPC), BPTES (GLS1), and Etomoxir (CPT1). **C.** ATP production by 8988-S and 8988-T cells measured by MitoStress test and MitoFuel Flex by Seahorse analysis. **D.** J_ATP_ quantification from glycolysis and OXPHOS. **E.** Examination of fold changes in metabolite pools related to glycolysis and citrate. **F.** Heatmap illustrating variations in glycan abundances influenced by glucose concentration in 8988-T cells. **G.** Flow cytometry analysis of glycan motifs between high and low glucose conditions for 8988-S and 8988-T cells. The plot shows capping motifs with a direct relationship between motif abundance and lectin/antibody binding. **H.** Summary elucidating the impact of metabolic differences between 8988-S and 8988-T cells on glycosylation. Asterisks with * indicating p < 0.05, ** for p < 0.01, *** for p < 0.001, and **** for p < 0.0001.

We asked whether the metabolite pools associated with glycolysis also showed differences in glucose metabolism (Figure 5E). Fructose-1,6-BP was significantly higher in the 8988-T cells, indicating a greater commitment of glucose to glycolysis, and intracellular lactate was higher with similar lactate secretion (Figure 5A), consistent with greater intracellular use of glucose. The total citrate pool (Figure 5E) and the ^13^C enrichment from glucose to citrate (Supplementary Figure 6) were also higher in the 8988-T cells, indicating a potentially greater glycolytic flux of carbons from glucose to the TCA cycle.

Therefore, these findings collectively support a model in which the 8988-T cells have greater glucose flux in glycolysis than 8988-S cells and greater glucose flux to non-glycosylation pathways such as the ribose production in the PPP, lactate production, the TCA cycle, and ATP production by glycolysis and OXPHOS. Despite higher glucose uptake and greater glucose flux in glycolysis, the 8988-T cells allocated less glucose to all monosaccharide production pathways except CMP-Neu5Ac.

The above findings also suggest that the increased reliance upon glucose for energy and non-glycosylation needs by the 8988-T cells results in low availability of glucose for glycosylation, and that the ability of the 8988-S cells to commit more glucose in glycosylation stems from its use of fatty acids for energy and the resultant availability of glucose for glycosylation. Thus, we asked whether changes in the amount of externally available glucose likewise modulated the amount of glucose committed to glycosylation. We cultured the 8988-T cells in a range of glucose concentrations for 5 days and quantified overall glycosylation using flow cytometry. The majority of glycans were substantially increased at glucose concentrations above 6 mM (Figure 5F), including N-glycans (Con-A), branching (PHA-L), and O-glycans (PNA and Jacalin). The decreased binding of certain lectins to single HexNAc moieties, such as VVL, LEL, and GSL II, is consistent with the exposure of short, truncated glycans upon the limitation of glycan extension. The 8988-S cells showed higher sensitivity to glucose concentration than the 8988T cells (Figure 5G), consistent with a lower requirement for glucose usage in non-glycosylation pathways.

These results support a model of regulation of glucose commitment glycosylation by glucose availability, where glucose demands in energy and non-glycosylation pathways determine glucose availability for glycosylation (Figure 5H). The 8988-T cells had higher glucose uptake but increased glucose flow to non-glycosylation pathways, which reduces the availability of fructose-6P for the nucleotide sugar synthesis essential to glycosylation (Figure 5H). The 8988-S cells, on the other hand, owing to their use of fatty acids for energy, required less glucose for non-glycosylation pathways and therefore had a higher glucose allocation to glycosylation.

## Discussion

Here, we report for the first time a novel method that combines stable isotope tracing with metabolomics to enable direct observations of glucose allocation to nucleotide sugars and cell-membrane glycans. Previous studies have shown associations between alterations in metabolism and altered glycosylation in glycoconjugates in cell membrane glycans. However, a direct link between glucose metabolism and altered glycan synthesis expression has not previously been established. We enabled unambiguous identification of isotope incorporation into cell membrane glycans by isolating monosaccharides from membrane-bound glycoconjugates and then resolving the monosaccharide isomers in high-performance liquid chromatography (HPLC) coupled with mass spectrometry. This method has overcome the challenge of distinguishing between multiple types of isomeric monosaccharides that have only minor structural differences. Here, we found that competing demands for glucose utilization for energy or other anabolic pathways can constrain glucose commitment to glycosylation. We have demonstrated that the synthesis of nucleotide sugars and cell membrane glycans was limited by the availability of glucose-6P and fructose-6P, rather than the quantity of glucose processed through glycolysis.

These findings suggest that metabolic shifts observed in cancer and immune cell activation alter nutrient flux to monosaccharide production and may therefore underlie the major changes observed in cell-membrane glycans. Such metabolic shifts occur, for example, in cancer EMT^37^ and the differentiation of naïve CD8+ T cells into effector T cells,^38,39^ both of which use more glucose to meet anabolic and energy requirements. Furthermore, steady-state differences in such metabolic phenotypes exist between T cells: cytotoxic and helper T cells primarily rely on aerobic glycolysis for their differentiation,^41,42^ whereas regulatory T and memory T cells predominantly utilize oxidative phosphorylation and fatty acid oxidation.^43,44^ Therefore, metabolic shifts that potentially affect glycosylation are prominent in multiple cellular processes. Our novel carbon tracing metabolomic method enables investigations into how the metabolic shifts affect cell-membrane glycosylation.

Our observation that altered glucose carbon usage regulates cell membrane glycan expression could further inform our understanding of tissue organization, cell surface receptor modulation, cellular immune responses, and cellular adhesion and migration. For example, increased complex N-glycosylation increases the retention, stability, clustering, and/or activation of cell-surface receptors.^45,46^ Increases in glycosphingolipids, including GM3 and SM4, help maintain KRAS plasma membrane localization,^47^ and changes in glycosylation of adhesion molecules like E-cadherin and integrins promote cell adhesion and migration.^48^ Elevations of sialic acid in cell-membrane glycans could contribute to cancer progression by dampening the immune response^49,50^ or could enable migration after epithelial-mesenchymal transition (EMT).^51,52^ In the immune system, changes in the glycosylation of membrane proteins and lipid rafts upon activation and differentiation of lymphocytes enable antigen recognition^53^ and homing of immune cells.^54–56^ Further investigations could show how metabolic shifts affect function through alterations in cell-membrane glycosylation. A key experiment then would be in vivo stable isotope tracing in mouse models of cancer,^8^ which would directly examine how altered cancer cell nutrient metabolism leads to altered membrane glycosylation and cancer progression.

The current study used cell lines derived from the same patient, which provided comparisons between genetically identical cells with differing metabolic requirements, but we do not yet know the generizability of the study to other settings. In addition, we only demonstrated differences in total amounts of monosaccharides rather than specific glycan motifs. The structures of glycans are regulated in part by the expression levels of ∼200 glycosyltransferases, many of which are redundant,^24^ and specific motifs are tightly controlled by key glycosyltransferases. For example, the ABO blood type glycans are controlled by the expression and genotype of the ABO glycosyltransferase,^57^ and the production of the Lewis X glycan comprising the CD15 epitope on granulocytes is controlled by the FUT4 and FUT9 glycosyltransferases.^58^ Further details could be probed using inhibitors of specific glycosylation pathways, such as N- and O-glycans and glycosphingolipids, or by presorting various types of intact glycans to localize isotope incorporation into specific features. Also, stable isotope detection and metabolomics on cell-surface glycans readily applies to other nutrients, such as glutamine, lactate, or galactose. The method presented here could be further developed for such studies.

In summary, we have developed a method for stable-isotope tracing and metabolomics to determine the commitment of media nutrients to cell-membrane glycans. The workflow enables studies of the connection between the flux of nutrients to non-glycosylation and glycosylation pathways and the effects of metabolic shifts upon cell-membrane glycosylation. In the system studied here, the commitment of glucose to membrane glycosylation was determined primarily by the availability of glucose relative to the use of glucose in non-glycosylation pathways. This relationship suggests a mode by which metabolism affects cell functions such signaling, immune recognition, and adhesion and migration.

## Acknowledgements

This work was supported by the Catalytic Fund for Metabolism Research at the Van Andel Institute. David M Brass PhD assisted in the preparation of this manuscript.

## References

1. Reily, C., Stewart, T. J., Renfrow, M. B. & Novak, J. Glycosylation in health and disease. Nat Rev Nephrol 15, 346–366 (2019).

2. Shen, X., Niu, N. & Xue, J. Oncogenic KRAS triggers metabolic reprogramming in pancreatic ductal adenocarcinoma. J. Transl. Intern. Med. 11, 322–329 (2022).

3. Li, J.-T., Wang, Y.-P., Yin, M. & Lei, Q.-Y. Metabolism remodeling in pancreatic ductal adenocarcinoma. Cell Stress 3, 361 (2019).

4. Araujo, L., Khim, P., Mkhikian, H., Mortales, C.-L. & Demetriou, M. Glycolysis and glutaminolysis cooperatively control T cell function by limiting metabolite supply to N-glycosylation. eLife 6, e21330 (2017).

5. Lucena, M. C. et al. Epithelial Mesenchymal Transition Induces Aberrant Glycosylation through Hexosamine Biosynthetic Pathway Activation*. J Biol Chem 291, 12917–12929 (2016).

6. Rossi, M. et al. PHGDH heterogeneity potentiates cancer cell dissemination and metastasis. Nature 605, 747–753 (2022).

7. Alisson-Silva, F. et al. Increase of O-Glycosylated Oncofetal Fibronectin in High Glucose-Induced Epithelial-Mesenchymal Transition of Cultured Human Epithelial Cells. PLoS ONE 8, e60471 (2013).

8. Ma, E. H. et al. Metabolic Profiling Using Stable Isotope Tracing Reveals Distinct Patterns of Glucose Utilization by Physiologically Activated CD8+ T Cells. Immunity 51, 856–870.e5 (2019).

9. Jang, C., Chen, L. & Rabinowitz, J. D. Metabolomics and Isotope Tracing. Cell 173, 822–837 (2018).

10. Sheldon, R. D., Ma, E. H., DeCamp, L. M., Williams, K. S. & Jones, R. G. Interrogating in vivo T-cell metabolism in mice using stable isotope labeling metabolomics and rapid cell sorting. Nat. Protoc. 16, 4494–4521 (2021).

11. Kim, J. et al. The hexosamine biosynthesis pathway is a targetable liability in KRAS/LKB1 mutant lung cancer. Nat Metabolism 2, 1401–1412 (2020).

12. Scherpenzeel, M. et al. Dynamic tracing of sugar metabolism reveals the mechanisms of action of synthetic sugar analogs. Glycobiology 32, cwab106 (2021).

13. Conte, F. et al. Isotopic Tracing of Nucleotide Sugar Metabolism in Human Pluripotent Stem Cells. Cells 12, 1765 (2023).

14. Trefely, S., Ashwell, P. & Snyder, N. W. FluxFix: automatic isotopologue normalization for metabolic tracer analysis. Bmc Bioinformatics 17, 485 (2016).

15. Mookerjee, S. A., Nicholls, D. G. & Brand, M. D. Determining Maximum Glycolytic Capacity Using Extracellular Flux Measurements. Plos One 11, e0152016 (2016).

16. Mookerjee, S. A., Gerencser, A. A., Nicholls, D. G. & Brand, M. D. Quantifying intracellular rates of glycolytic and oxidative ATP production and consumption using extracellular flux measurements. J Biol Chem 292, 7189–7207 (2017).

17. Cebin, A. V., Komes, D. & Ralet, M.-C. Development and Validation of HPLC-DAD Method with Pre-Column PMP Derivatization for Monomeric Profile Analysis of Polysaccharides from Agro-Industrial Wastes. Polymers-basel 14, 544 (2022).

18. Wang, W. et al. Optimization of reactions between reducing sugars and 1-phenyl-3-methyl-5-pyrazolone (PMP) by response surface methodology. Food Chem 254, 158–164 (2018).

19. Xu, G., Amicucci, M. J., Cheng, Z., Galermo, A. G. & Lebrilla, C. B. Revisiting monosaccharide analysis – quantitation of a comprehensive set of monosaccharides using dynamic multiple reaction monitoring. Analyst 143, 200–207 (2017).

20. Honda, S. et al. High-performance liquid chromatography of reducing carbohydrates as strongly ultraviolet-absorbing and electrochemically sensitive 1-phenyl-3-methyl5-pyrazolone derivatives. Anal. Biochem. 180, 351–357 (1989).

21. Reily, C., Stewart, T. J., Renfrow, M. B. & Novak, J. Glycosylation in health and disease. Nat Rev Nephrol 15, 346–366 (2019).

22. Harazono, A. et al. A comparative study of monosaccharide composition analysis as a carbohydrate test for biopharmaceuticals. Biologicals 39, 171–180 (2011).

23. Kim, S., Kim, S. I., Ha, K.-S. & Leem, S.-H. An improved method for quantitative sugar analysis of glycoproteins. Exp Mol Medicine 32, 141–145 (2000).

24. Schjoldager, K. T., Narimatsu, Y., Joshi, H. J. & Clausen, H. Global view of human protein glycosylation pathways and functions. Nature Reviews Molecular Cell Biology 21, (2020).

25. Klamer, Z., Hsueh, P., Ayala-Talavera, D. & Haab, B. Deciphering Protein Glycosylation by Computational Integration of On-chip Profiling, Glycan-array Data, and Mass Spectrometry* [S]. Mol Cell Proteomics 18, 28–40 (2019).

26. Nohara, K., Wang, F. & Spiegel, S. Glycosphingolipid composition of MDA-MB-231 and MCF-7 human breast cancer cell lines. Breast Cancer Res. Treat. 48, 149–157 (1998).

27. Cui, H., Lin, Y., Yue, L., Zhao, X. & Liu, J. Differential expression of the α2,3-sialic acid residues in breast cancer is associated with metastatic potential. Oncol. Rep. 25, 1365–71 (2011).

28. Pally, D. et al. Heterogeneity in 2,6-Linked Sialic Acids Potentiates Invasion of Breast Cancer Epithelia. Acs Central Sci 7, 110–125 (2021).

29. Rodriguez, E. et al. Analysis of the glyco-code in pancreatic ductal adenocarcinoma identifies glycan-mediated immune regulatory circuits. Commun Biology 5, 41 (2022).

30. Cumin, C. et al. Glycosphingolipids are mediators of cancer plasticity through independent signaling pathways. Cell Reports 40, 111181 (2022).

31. Kim, P. K. et al. Hyaluronic acid fuels pancreatic cancer cell growth. Elife 10, e62645 (2021).

32. Akella, N. M., Ciraku, L. & Reginato, M. J. Fueling the fire: emerging role of the hexosamine biosynthetic pathway in cancer. BMC Biol. 17, 52 (2019).

33. Kim, P. K. et al. Hyaluronic Acid Fuels Pancreatic Cancer Growth. Biorxiv 2020.09.14.293803 (2020) doi:10.1101/2020.09.14.293803.

34. Daemen, A. et al. Metabolite profiling stratifies pancreatic ductal adenocarcinomas into subtypes with distinct sensitivities to metabolic inhibitors. Proc National Acad Sci 112, E4410– E4417 (2015).

35. Holst, S., Belo, A. I., Giovannetti, E., Die, I. van & Wuhrer, M. Profiling of different pancreatic cancer cells used as models for metastatic behaviour shows large variation in their N-glycosylation. Sci Rep-uk 7, 16623 (2017).

36. Zhang, T. et al. Differential O- and Glycosphingolipid Glycosylation in Human Pancreatic Adenocarcinoma Cells With Opposite Morphology and Metastatic Behavior. Front. Oncol. 10, 732 (2020).

37. Jia, D. et al. Towards decoding the coupled decision-making of metabolism and epithelial-to-mesenchymal transition in cancer. Br. J. Cancer 124, 1902–1911 (2021).

38. Pearce, E. L., Poffenberger, M. C., Chang, C.-H. & Jones, R. G. Fueling Immunity: Insights into Metabolism and Lymphocyte Function. Science 342, 1242454 (2013).

39. Ma, E. H. et al. Metabolic Profiling Using Stable Isotope Tracing Reveals Distinct Patterns of Glucose Utilization by Physiologically Activated CD8+ T Cells. Immunity 51, 856–870.e5 (2019).

40. Spelat, R. et al. Metabolic reprogramming and membrane glycan remodeling as potential drivers of zebrafish heart regeneration. *Commun*. Biol. 5, 1365 (2022).

41. Gubser, P. M. et al. Rapid effector function of memory CD8+ T cells requires an immediate-early glycolytic switch. Nat. Immunol. 14, 1064–1072 (2013).

42. Klysz, D. et al. Glutamine-dependent α-ketoglutarate production regulates the balance between T helper 1 cell and regulatory T cell generation. Sci. Signal. 8, ra97 (2015).

43. Michalek, R. D. et al. Cutting Edge: Distinct Glycolytic and Lipid Oxidative Metabolic Programs Are Essential for Effector and Regulatory CD4+ T Cell Subsets. J. Immunol. 186, 3299–3303 (2011).

44. Shi, L. Z. et al. HIF1α–dependent glycolytic pathway orchestrates a metabolic checkpoint for the differentiation of TH17 and Treg cells. J. Exp. Med. 208, 1367–1376 (2011).

45. Zhou, Q. & Qiu, H. The Mechanistic Impact of N-Glycosylation on Stability, Pharmacokinetics, and Immunogenicity of Therapeutic Proteins. J. Pharm. Sci. 108, 1366–1377 (2019).

46. GC, S., Bellis, S. L. & Hjelmeland, A. B. ST6Gal1: Oncogenic signaling pathways and targets. Frontiers Mol Biosci 9, 962908 (2022).

47. Liu, J. et al. Glycolysis regulates KRAS plasma membrane localization and function through defined glycosphingolipids. Nat. Commun. 14, 465 (2023).

48. Bassagañas, S. et al. Pancreatic Cancer Cell Glycosylation Regulates Cell Adhesion and Invasion through the Modulation of α2β1 Integrin and E-Cadherin Function. Plos One 9, e98595 (2014).

49. Cai, X., Thinn, A. M. M., Wang, Z., Shan, H. & Zhu, J. The importance of N-glycosylation on β3 integrin ligand binding and conformational regulation. Sci. Rep. 7, 4656 (2017).

50. Cagnoni, A. J., Sáez, J. M. P., Rabinovich, G. A. & Mariño, K. V. Turning-Off Signaling by Siglecs, Selectins, and Galectins: Chemical Inhibition of Glycan-Dependent Interactions in Cancer. Front. Oncol. 6, 109 (2016).

51. Beaman, E.-M., Carter, D. R. F. & Brooks, S. A. GALNTs: master regulators of metastasis-associated epithelial-mesenchymal transition (EMT)? Glycobiology 32, 556–579 (2022).

52. Cumin, C. et al. Glycosphingolipids are mediators of cancer plasticity through independent signaling pathways. Cell Rep. 40, 111181 (2022).

53. Comelli, E. M. et al. Activation of Murine CD4+ and CD8+ T Lymphocytes Leads to Dramatic Remodeling of N-Linked Glycans. J. Immunol. 177, 2431–2440 (2006).

54. Zeng, J. et al. Cosmc controls B cell homing. Nat. Commun. 11, 3990 (2020).

55. Nagafuku, M., et al. CD4 and CD8 T cells require different membrane gangliosides for activation. Proc. Natl. Acad. Sci. 109, E336–E342 (2012).

56. Togayachi, A. et al. Polylactosamine on glycoproteins influences basal levels of lymphocyte and macrophage activation. Proc. Natl. Acad. Sci. 104, 15829–15834 (2007).

57. Yamamoto, F., Clausen, H., White, T., Marken, J. & Hakomori, S. Molecular genetic basis of the histo-blood group ABO system. Nature 345, 229–233 (1990).

58. Nakayama, F. et al. CD15 Expression in Mature Granulocytes Is Determined by α1,3-Fucosyltransferase IX, but in Promyelocytes and Monocytes by α1,3-Fucosyltransferase IV*. J. Biol. Chem. 276, 16100–16106 (2001).

