## Supplementary Figures for "Metabolomics and ^13^C Labelled Glucose Tracing to Identify Carbon Incorporation into Aberrant Cell Membrane Glycans in Cancer"

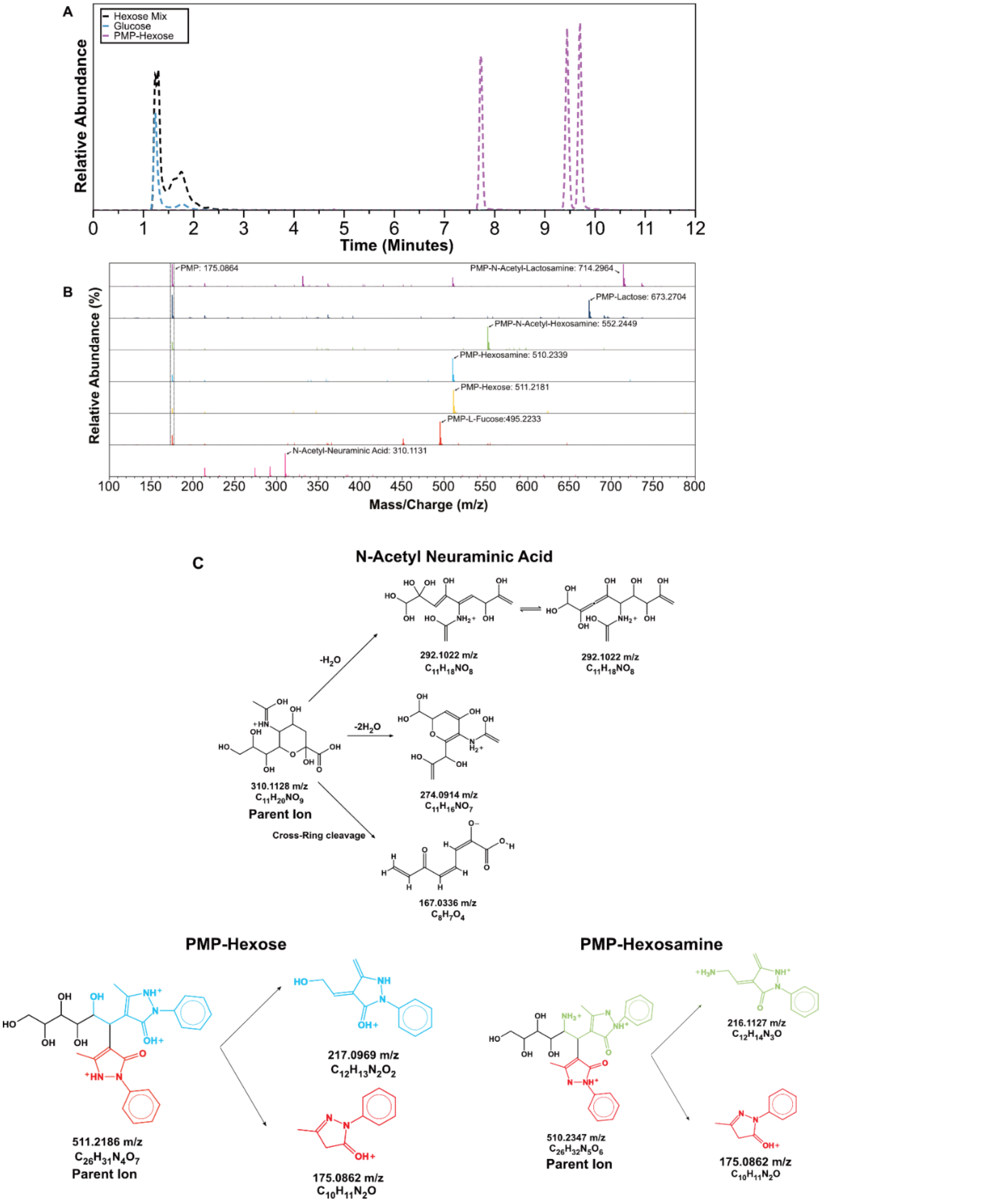


**Supplementary Figure 1. A.** Chromatogram of hexoses shown before and after PMP derivatization. **B.** MS1 spectra of PMP-derivatized carbohydrates with parental ions. **C.** MS2 spectra fragments of standard carbohydrates derivatized with PMP (except Neu5Ac).

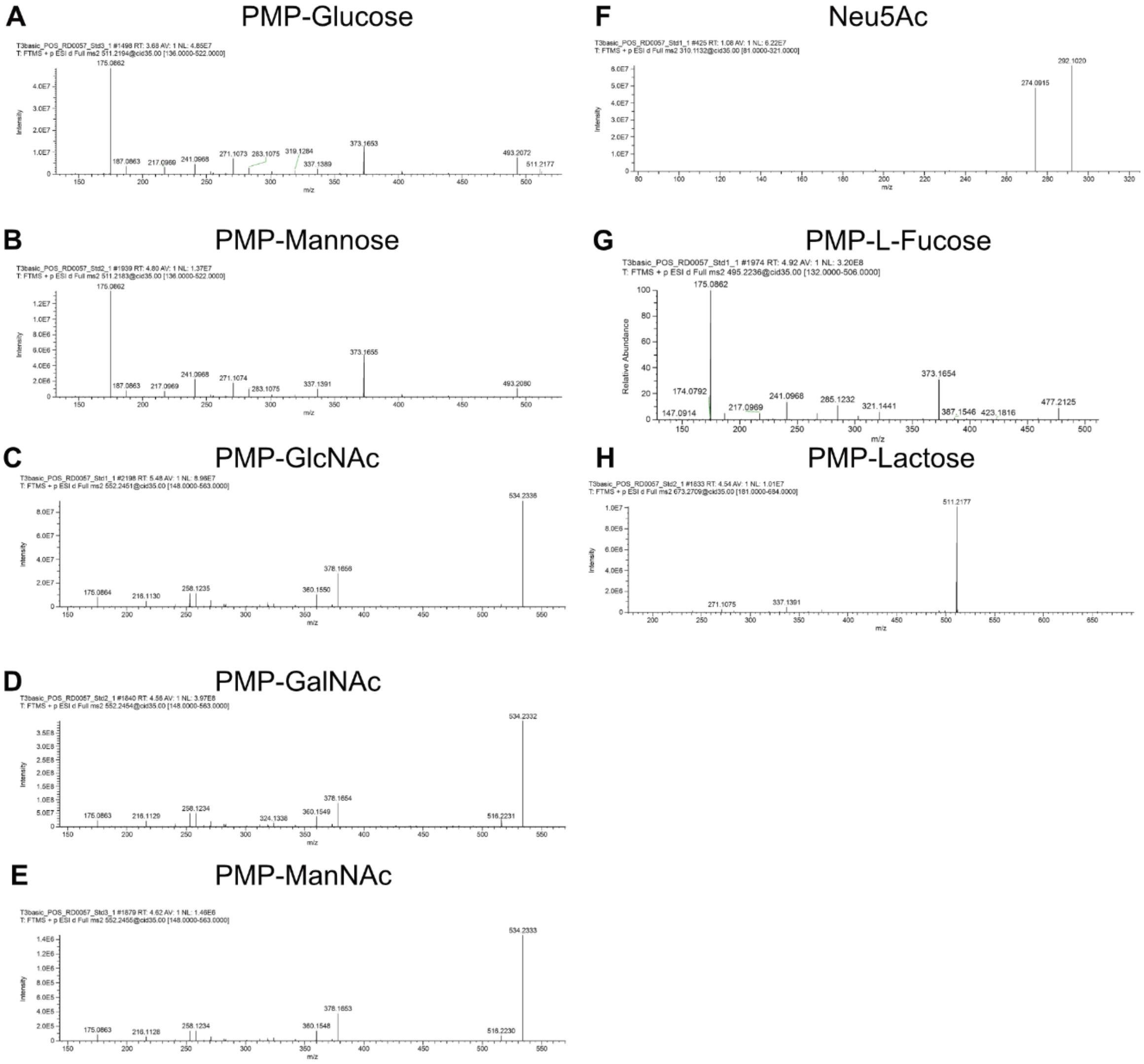


**Supplementary Figure 2.** CID MS2 spectra for standards used in LC-MS method.

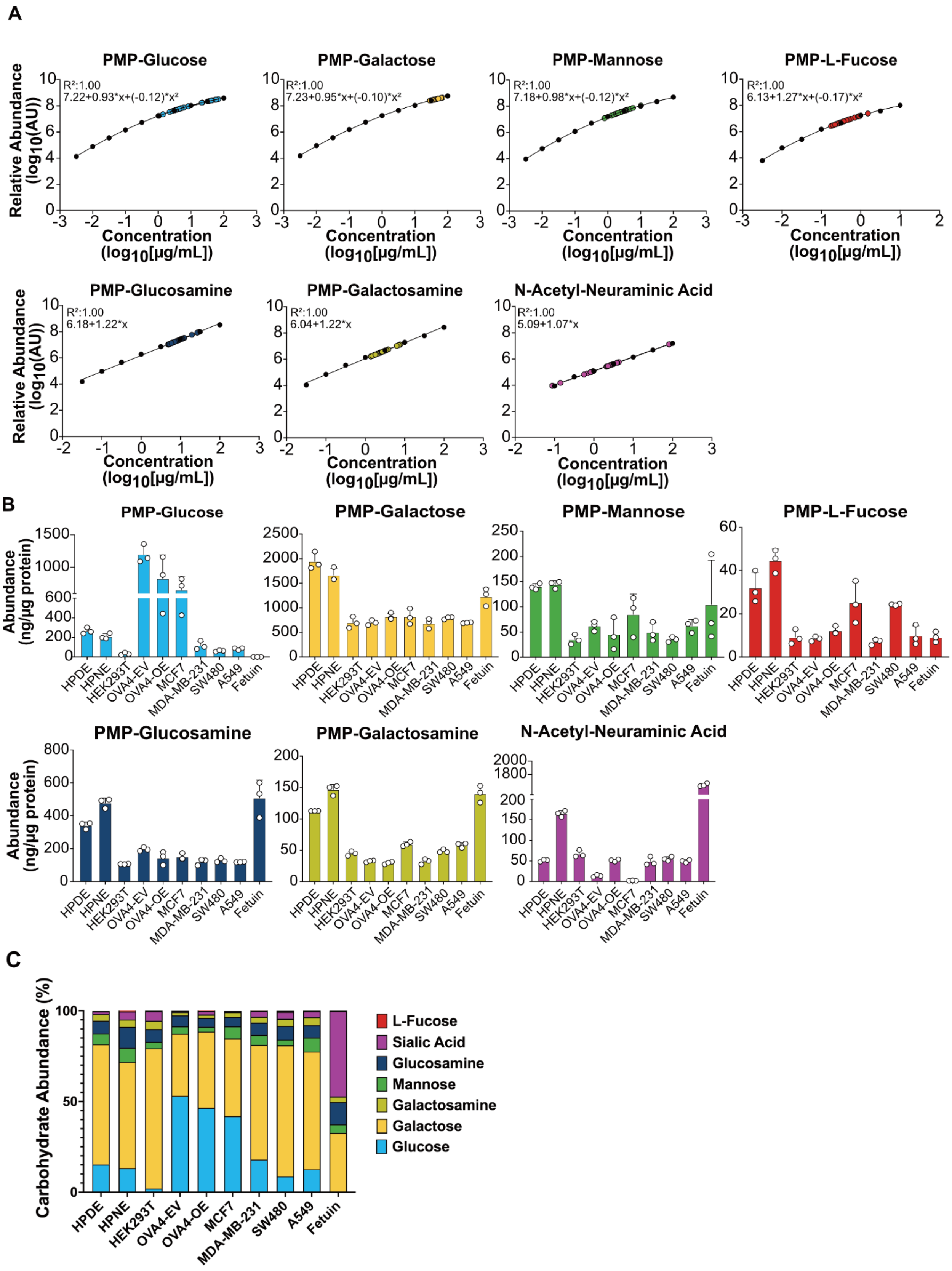
A
PMP-Glucose
R²:1.00
7.22+0.93*x+(-0.12)*x²
PMP-Galactose
R²:1.00
7.23+0.95*x+(-0.10)*x²
PMP-Mannose
R²:1.00
7.18+0.98*x+(-0.12)*x²
PMP-L-Fucose
R²:1.00
6.13+1.27*x+(-0.17)*x²
PMP-Glucosamine
R²:1.00
6.18+1.22*x
PMP-Galactosamine
R²:1.00
6.04+1.22*x
N-Acetyl-Neuraminic Acid
R²:1.00
5.09+1.07*x
Relative Abundance (log10(AU))
Concentration (log10[μg/mL])
B
Abundance (ng/μg protein)
PMP-Glucose
PMP-Galactose
PMP-Mannose
PMP-L-Fucose
PMP-Glucosamine
PMP-Galactosamine
N-Acetyl-Neuraminic Acid
HPDE
HPNE
HEK293T
OVA4-EV
OVA4-OE
MCF7
MDA-MB-231
SW480
A549
Fetuin
C
Carbohydrate Abundance (%)
L-Fucose
Sialic Acid
Glucosamine
Mannose
Galactosamine
Galactose
Glucose

**Supplementary Figure 3. A.** External standard curves for monosaccharides detected in samples. PMP-hexoses and PMP-L-fucose were fitted to a quadratic nonlinear regression due to the high dynamic range, and PMP-hexosamines and N-acetyl-neuraminic acid were fitted to linear regression. **B.** Absolute quantification of 7 monosaccharides across different cell lines and fetuin, a glycoprotein used as control. Samples were normalized by protein content. **C.** Percentage of monosaccharide distribution per sample.

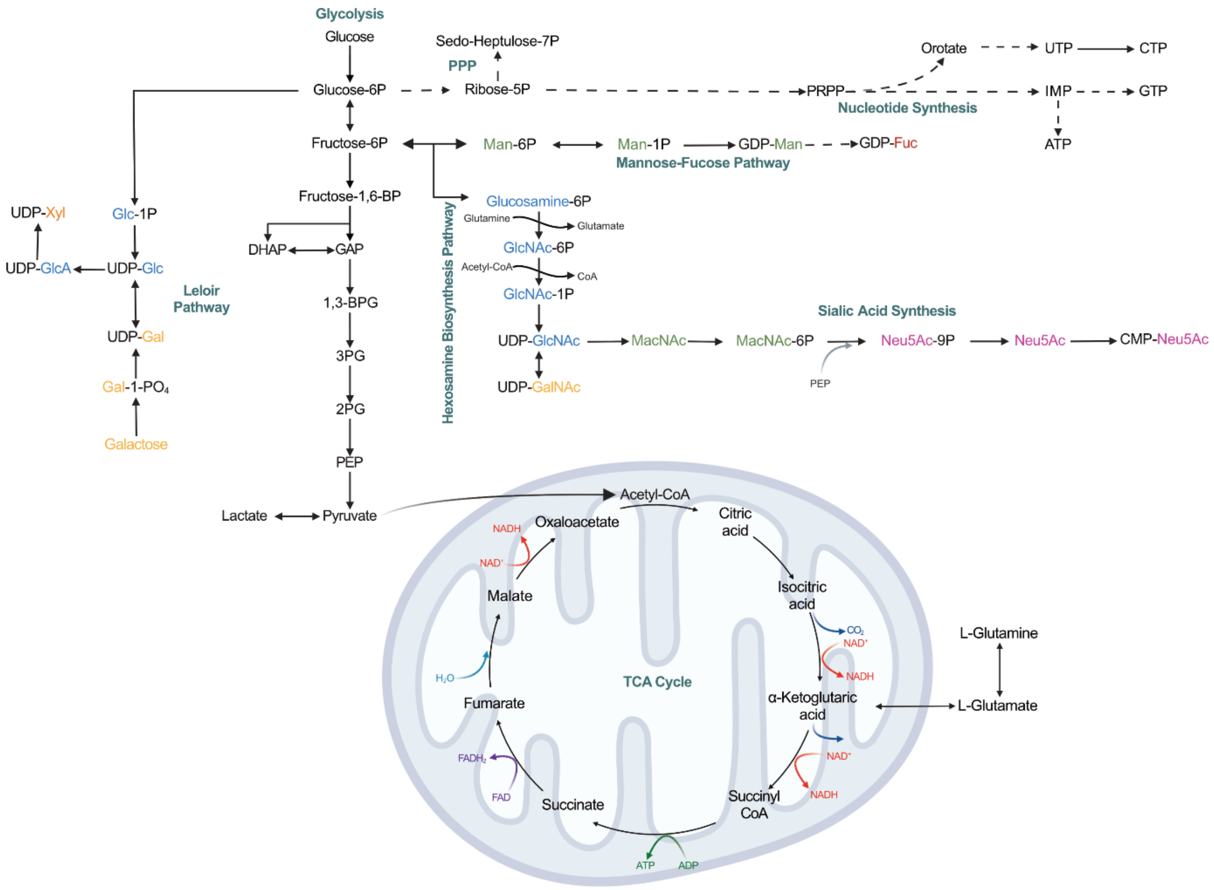


**Supplementary Figure 4. A.** Diagram representing central carbon metabolism and offshoot pathways related to nucleotide sugar synthesis. Glyceraldehyde 3-Phosphate (GAP), Dihydroxy-acetone-Phosphate (DHAP), 1,3-Bisphophosglycerate (1,3-BPG), 3-Phosphoglycerate (3PG), 2-Phosphoglycerate (2PG), Phosphoenolpyruvate (PEP). Pathways are represented with bold green text.

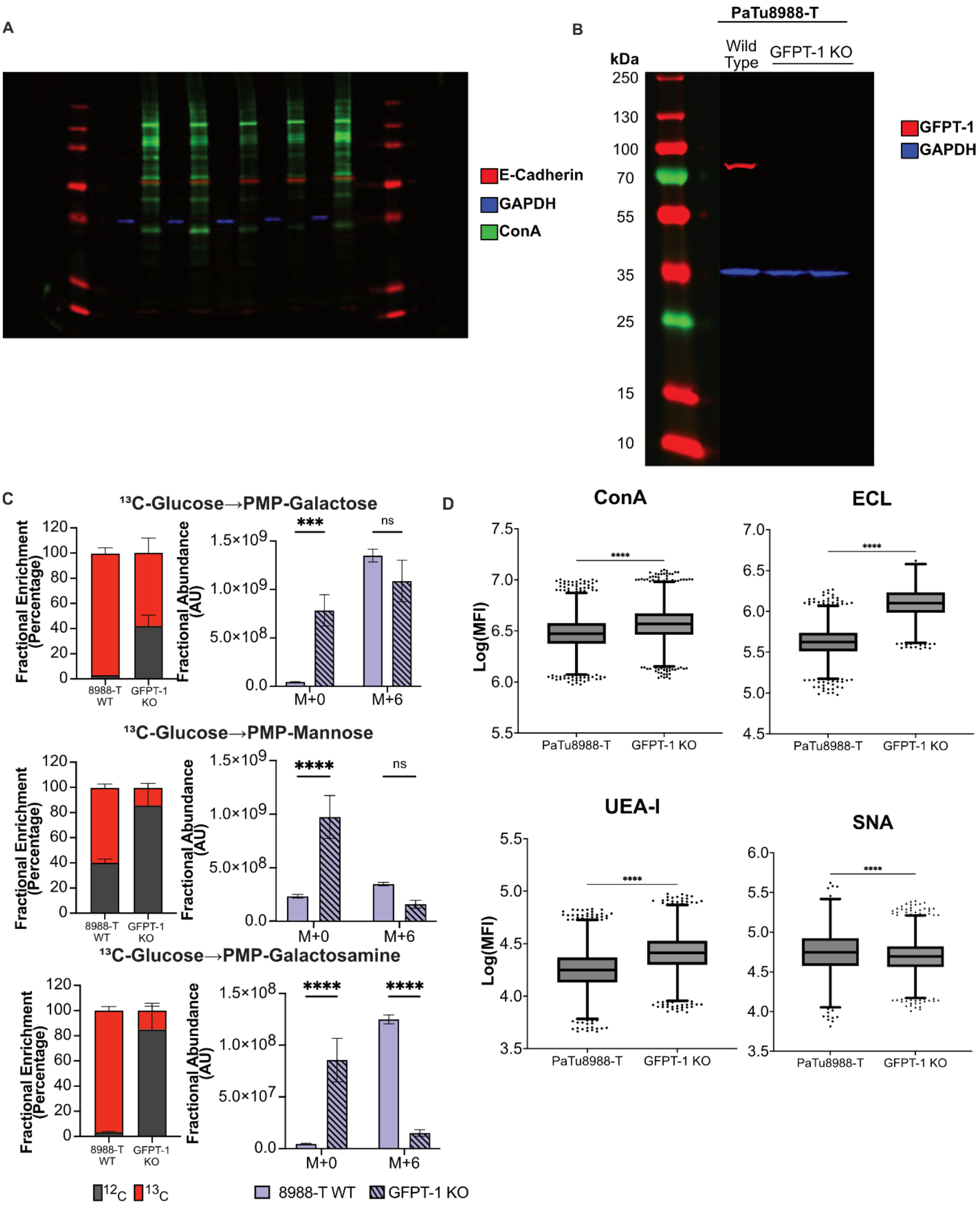


**Supplementary Figure 5. A.** Western blot of cytosolic and membrane fractions. Red (E-Cadherin), Blue (GAPDH), Green (Con-A). **B.** Western blot of GFPT-1 KO cell lines. **C.** Relative abundance of PMP-Mannose and PMP-galactose in 8988-T WT and GFPT-1 KO cells **D.** Flow cytometry analysis between 8988-T WT and GFPT-1 KO cells showing N-Glycans (ConA),

Galactose-GlcNAc (ECL), Fucoseα1-2Galβ1-4GlcNAc (UEA I), and α 2-6-sialylated LacNAc (SNA). Each dot represents a cell event.

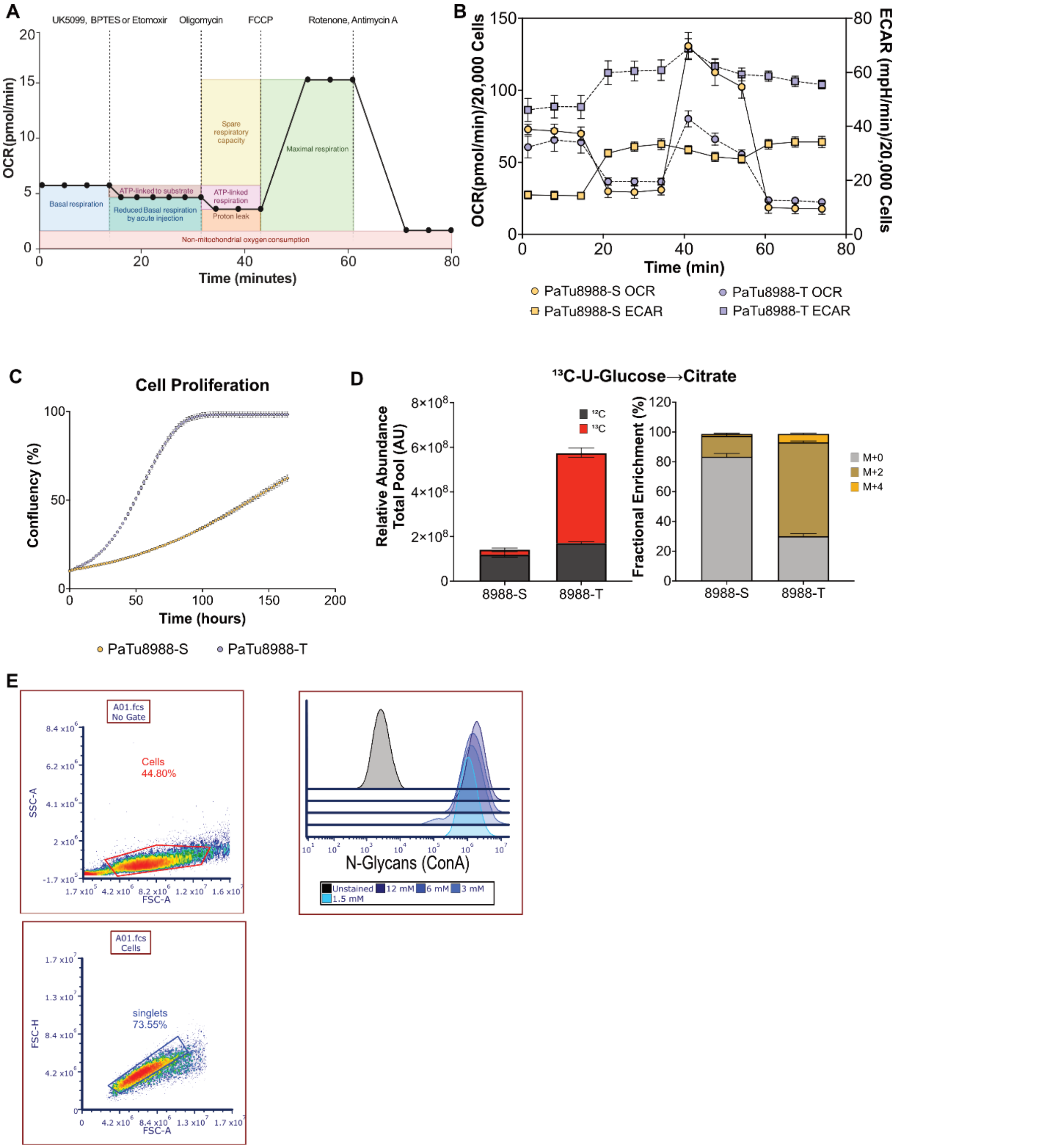


**Supplementary Figure 6. A.** Seahorse analysis strategy used for acute inhibition of MPC (UK5099), GLS1 (BPTES), and Etomoxir (CPT1A) to measure nutrient dependency on glucose, L-glutamine, and fatty acids, respectively. Oligomycin inhibits ATPase, FCCP is a membrane decoupler, and Rotenone and antimycin A inhibit complexes 1 and 3 of the ETC, respectively. **B.** SeaHorse analysis using MitoStress Test showing OCR and ECAR used to calculate $J_{ATP}$. **C.**

Proliferation rate of 8988-S and 8988-T cells. **D.** Total citrate pool and MID. **E.** Gating strategy used in flow cytometry.
